# Variable bites and dynamic populations; new insights in *Leishmania* transmission

**DOI:** 10.1101/2020.09.07.285809

**Authors:** Samuel Carmichael, Ben Powell, Thomas Hoare, Pegine Walrad, Jon Pitchford

## Abstract

Leishmaniasis is a neglected tropical disease which kills an estimated 50000 people each year, with its deadly impact confined mainly to lower to middle income countries. *Leishmania* parasites are transmitted to human hosts by sand fly vectors during blood feeding. Recent experimental work shows that transmission is modulated by the patchy landscape of infection in the host’s skin, and the parasite population dynamics within the vector. Here we assimilate these new findings into a simple probabilistic model for disease transmission which replicates recent experimental results, and assesses their relative importance. The results of subsequent simulations, describing random parasite uptake and dynamics across multiple blood meals, show that skin heterogeneity is important for transmission by short-lived flies, but that for longer-lived flies with multiple bites the population dynamics within the vector dominate transmission probability. Our results indicate that efforts to reduce fly lifespan beneath a threshold of around two weeks may be especially helpful in reducing disease transmission.

## Introduction

Leishmaniasis is caused by parasites of the *Leishmania* genus. Details of the infection depend on the particular species [1], but all species share the same general vector-borne lifecycle, with distinct and complex life cycle stages in the mammalian host and sand fly vector [2]. *Leishmania* parasites have two main morphological forms. Broadly speaking, Amastigotes (ovoid, non-flagellated) dominate the mammalian stage of the lifecycle. Promastigotes (larger, flagellated) are found in the vector, and are divided into multiple developmental subclasses [3, 4].

When an uninfected female sand fly bites an infected mammal, it ingests amastigote infected macrophages from the host’s skin or blood [5]. Amastigotes differentiate into procyclic promastigotes, which are resistant to the digestive enzymes of the sand fly midgut [2]. Procyclics then replicate before differentiating into nectomonad promastigotes [3]. Nectomonads are able to migrate towards the thoracic midgut [2] and bind to the midgut epithelium [6]. There they differentiate into leptomonad promastigotes [3]. Leptomonads are the final replicative stage, replicating more rapidly than procyclics, and migrating through the thoracic midgut to the stomodeal valve [3]. Finally, leptomonads differentiate into metacyclic promastigotes, the infectious stage. Metacyclics have a short cell body and long flagellum to enhance motility [3]; they can be transmitted to a new host where they infect host macrophages via phagocytosis. (The infection dynamics in the host are similarly complex [7] [8], but are not relevant to this investigation which focuses on transmission from vector to host.)

Two recent key findings concerning details of Leishmania biology offer new insights into the possibility of understanding, and possibly controlling, the spread of the disease. They are described below:

### Patchy landscape of infection in the host

Transmission from host to vector occurs when a sand fly consumes a blood meal from an infected host. Doehl *et al*. [5] showed first that the parasite load in the host’s skin, rather than that in its blood, is the major determinant of successful infection, and furthermore that skin parasite burden is highly variable within and between hosts. They then used a modelling approach to investigate the consequences of this patchiness. For a host with a low mean parasite burden, a patchy skin landscape enhanced outward transmission (although the overall probability of successful transmission remained low), whereas for a host with a high parasite burden a homogenous distribution favoured transmission.

### Retroleptomonads

A new lifecycle stage was identified by Serafim et al. [9], the retroleptomonad promastigote [9]. When an infected sand fly takes another blood meal, the metacyclic stage can de-differentiate into a leptomonad-like stage, termed the retroleptomonad. These replicate for 3-4 days before differentiating back into metacyclics [9]. This serves to greatly amplify the parasite load prior to the next bite (4.5 fold increase in the number of metacyclics 18 days post infection in comparison to a single bite sand fly) and thus increases the probability of disease transmission [9], a finding confirmed experimentally under laboratory conditions.

The objective of this study was to build a mathematical model to incorporate these new findings, and thereby to assess how they might influence our understanding of the factors governing *Leishmania* transmission. A simple differential equation model, parameterised by data from [3], was developed to describe the population dynamics of nectomonad, leptomonad and metacyclic promastigote stages within the vector (Model A). This model was then refined by the addition of the retroleptomonad lifecycle stage, using data and observations from [9] (Model B). These models of population dynamics within the sand fly provide a framework for a series of stochastic simulations which describe the random processes of feeding and parasite ingestion across multiple blood meals. Such simulations allow the consequences of changes in disease prevalence at the epidemiological scale and the thresholds of disease transmission to be quantifiably predicted.

## 1 Model Details

### 1.1 Modelling Approach

The modelling strategy is summarised in Fig 1. First, we develop a simple, algebraically tractable and computationally efficient model for parasite population dynamics within a single infected sand fly, and then parameterise this model according to the available information. This model then forms a key ingredient in a series of larger stochastic simulations intended to extract useful details about the transmission of *Leishmania*.

**Fig 1.**
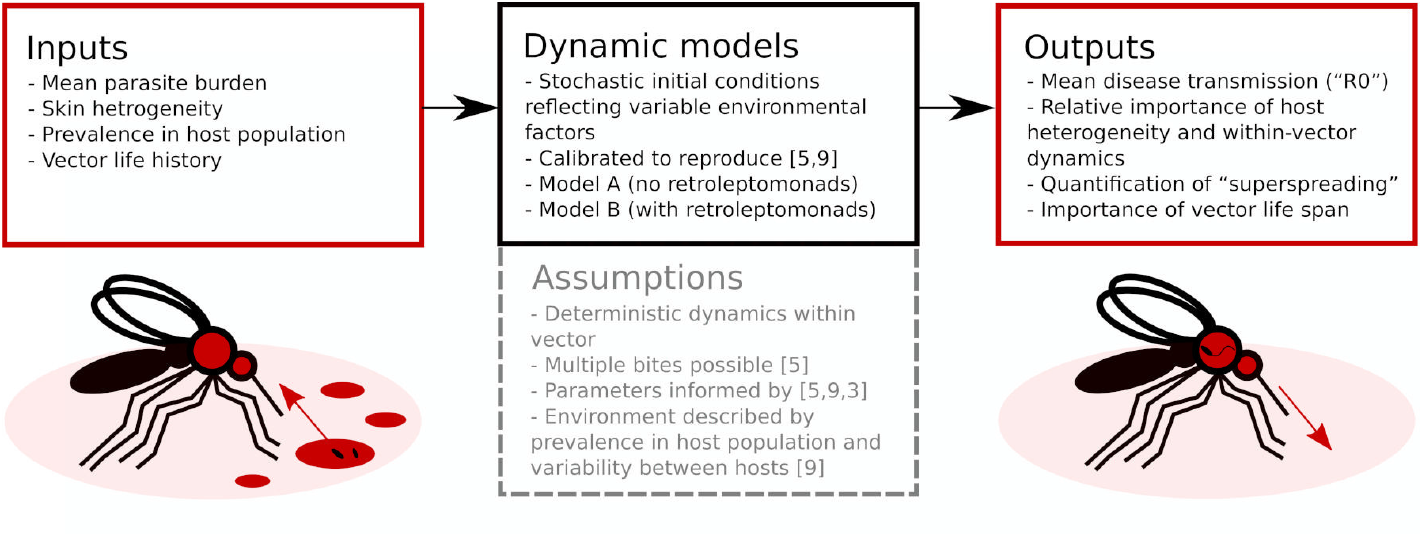
Flowchart overview of the modelling approach. Two dynamic models, calibrated to replicate prior results, evaluate parasite population dynamics in the sandfly vector. These can be used as part of larger simulations to obtain insights into *Leishmania* transmission.

In order to create a tractable model, several key assumptions are made. In addition to those represented in Fig 1, we also assume that differentiation between parasite life stages occurs at 100% efficiency and that there is a single globally applied carrying capacity.

### 1.2 Model Definitions

Model A describes the dynamics of Nectomonads (*N*), Leptomonads (*L*) and Metacyclics (*M*) using a simple set of near-linear ordinary differential equations (ODEs),

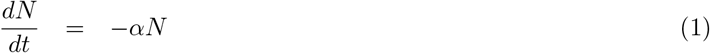

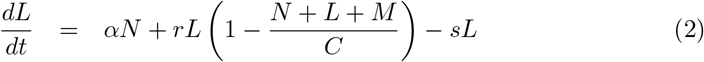

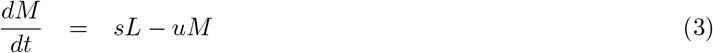

The assumptions are biologically parsimonious: *N* differentiate into *L* at rate *α, L* replicate at rate *r* (limited by a carrying capacity *C*) and differentiate to *M* at rate *s*, and *M* are also subject to mortality at rate *u*.

Model B extends Model A to incorporate the dynamics of the Retroleptomonads (*R*) [9] using two sets of near-linear ODEs. Under standard conditions ‘normal mode’ is used,

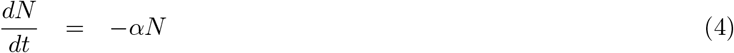

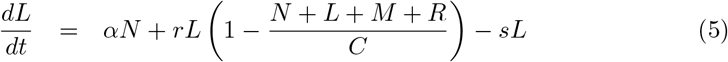

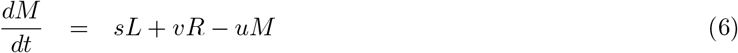

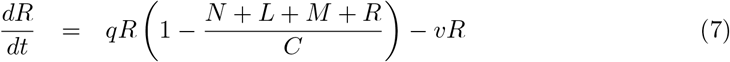

In addition to the original assumptions, it is assumed that any existing *R* differentiate to *M* at rate *v* and replicate at rate *q* limited by carry capacity *C*. For a four-day period after subsequent bites ‘dedifferentiation mode’ is used,

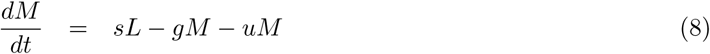

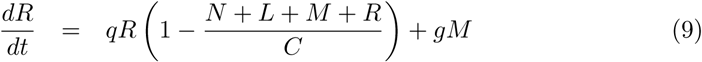

Now, *M* dedifferentiate to *R* at rate *g* and *R* no longer differentiate to *M*.

Parameterisation of Model A was performed using data obtained from Rogers *et al* [3] (see supplementary method S1) but due to a lack of suitable data, it was not possible to perform similar parameter fitting for the new parameters in Model B. Table 1 includes a summary of the default parameter values chosen.

**Table 1.** Table of default model parameter values.

| Parameter | Name | Default Value | Source |
| --- | --- | --- | --- |
| $\alpha$ | Nectomonad differentiation rate | 1.52 | [A] |
| $r$ | Leptomonad replication rate | 1.45 | [A] |
| $s$ | Leptomonad differentiation rate | 1.65 | [A] |
| $u$ | Metacyclic decline rate | 1.61 | [A] |
| $C$ | Carry capacity | $2 * 10^6$ | [A] |
| $v$ | Retroleptomonad differentiation rate | 4.0 | [B] |
| $q$ | Retroleptomonad replication rate | 3.5 | [B] |
| $g$ | Metacyclic dedifferentiation rate | 4.0 | [B] |

All parameters and their default values. [A]: Values are derived from parameterisation based on data from Rogers *et al*. [3], see supplementary method S1. [B]: Values chosen as a result of personal communications with Dr. Pegine Walrad.

## Results

### 1.2.1 Replicating experimental results on sand fly feeding schedules and mammalian infection heterogeneity

In order to verify that our retroleptomonad-inclusive Model B is capable of replicating the experimental results observed by Serafim *et al*. [9], we ran a set of 20000 Monte Carlo simulations designed to imitate their experimental setup. In this scenario, all flies take a bite at day 0 from an infected host. Half the flies take an additional bite at day 12 from an uninfected host, the other half take no subsequent bites. We fix the mean skin parasite burden to 2 × 10^6^ and let *k* = 2 to mimic the blood source used by Serafim *et al*.. After the initial bite, we take up a number of amastigotes according to the methods in supplementary methods S2. In this example, the initial number of nectomonads N_0_ has mean *µ* and variance *σ*^2^:

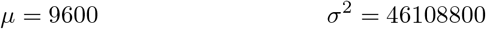

Of particular interest are the numbers of metacyclics and retroleptomonads present in each fly throughout their adult lifespan. Fig 2A compares the numbers of metacyclics and retroleptomonads at each day sampled by Serafim *et al*.

**Fig 2.**
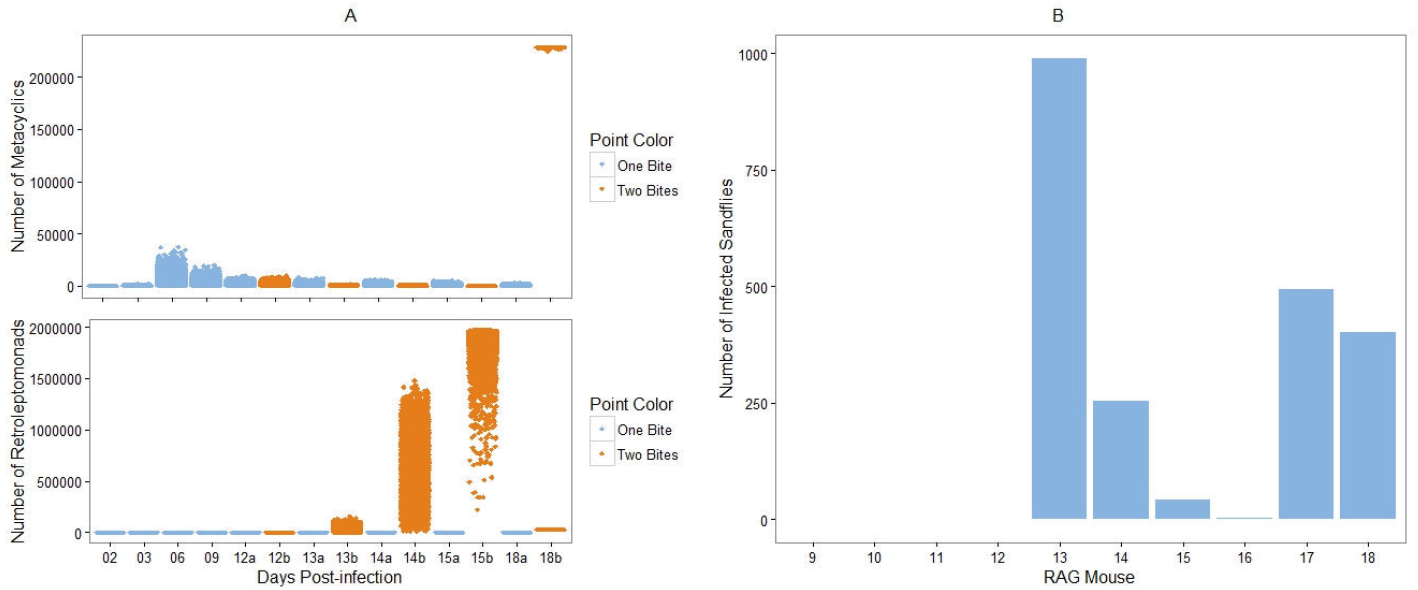
**Replicating the results of [9] and [5].** A) Comparison of the numbers of metacyclics (top) and retroleptomonads (bottom) at specific days throughout the lifespan of the simulated flies. Blue represents flies that bite only at day 0, orange represents flies that bite at day 12. The two categories are combined prior to day 12. B) Number of simulated sandflies considered infected at 7 days post-infection for RAG mice 10-18, parameterised according to Doehl *et al*.

Fig 2A reflects the qualitative dynamics observed in the experiments of Serafim *et al*. We observe a similar reduction in the number of metacyclics immediately after the bite at day 12 and a corresponding increase in the number of retroleptomonads over the same time period. Similar behaviour can be observed for the proportions of metacyclics and retroleptomonads (Supplementary Figure 1), and this behaviour is sufficiently robust to be observed even with parameter randomisation (Supplementary Figure 2).

We also wish to verify that our model can capture the importance of micro- and macro-scale heterogeneity in the skin parasite distribution as reported by Doehl *et al* [5]. To do so, we ran sets of 1000 Monte Carlo simulations for parameter combinations corresponding to mice 10-18 as calculated by Doehl *et al*. Each simulated fly bit an infected host at *t* = 0. We then sampled the number of metacyclics in each fly after 7 days, and calculated how many sandflies were infected at that time. For a sandfly to be considered infected, 500 metacyclics must be present. Fig 2B compares the number of infected sandflies for each mouse.

We observe that heavily infected mice such as mouse 13 are able to infect a large proportion, if not all, of the sandflies. Lighter infections such as those of mice 10 and 16 typically infect very few sandflies, if any. This matches the observations made by Doehl *et al* [5] and verifies that our model successfully captures the relationship between outward transmission potential and micro- and macro-scale skin patchiness.

### 1.3 Analytic results

In this section we provide a selection of analytically-derived properties and consequences of our proposed models. They serve to reinforce and validate the numerically derived behaviours discussed in Section 1.4. In particular, we present expressions bounding implied disease transmission probabilities in a range of hypothetical scenarios.

Specifically, we restrict our attention to scenarios in which a sand fly makes either two or three bites over a period of 12 days. In all scenarios the fly is assumed to bite an infected host at time *t* = 0, when it ingests *N*_0_ parasites in the nectomonad life stage, and an uninfected host at time *t* = 12, when it deposits *M*_12_ parasites in the metacyclic life stage. *N*_0_ is considered a random variable. *M*_12_ is considered a deterministic function of *N*_0_, so inherits probabilistic behaviour from this random variable. A transmission event is associated with the sand fly depositing a number of parasites exceeding a threshold *T*. Thus transmission is also a random variable inheriting probabilistic behaviours from *N*_0_.

The scenarios we consider differ in terms of the occurrence of an additional bite of an infected host at time *t* = 6. In our model the blood meal ingested in this bite triggers the retroleptomonad reproduction mechanism, effecting the number of metacyclics that can be deposited at time *t* = 12.

The structure of the model described in Section 1 is such that, given that there are bites only at times 0 and 12, *M*_12_ is in fact proportional to *N*_0_ i.e.

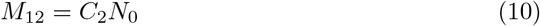

where *C*_2_ is a constant derived by solving the system of equations in Section 1. It is implicitly a function of the model’s differentiation rate parameters and the time elapsed between bites.

If the blood meal bite at time *t* = 6 does take place a different set of equations, involving the retroleptomonads, is used to determine the resulting number of metacyclics at time *t* = 12. *M*_12_ is now determined by *N*_0_ and a correspondingly different multiplicative constant

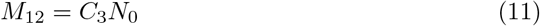

Expressions (10) and (11) can be combined when we write

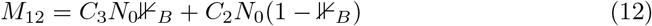

where 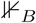 is an indicator function taking value one when the blood meal bite at time *t* = 6 does take place and zero otherwise.

We can now, for instance, consider the expectation of *M*_12_

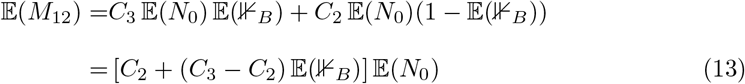

which follows on the assumption that 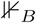 and *N*_0_ are considered probabilistically independent. Note that 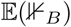 is the probability that the blood meal bite takes place.

Eq (12) can also be used to produce an expression for the transmission probability at time *t* = 12

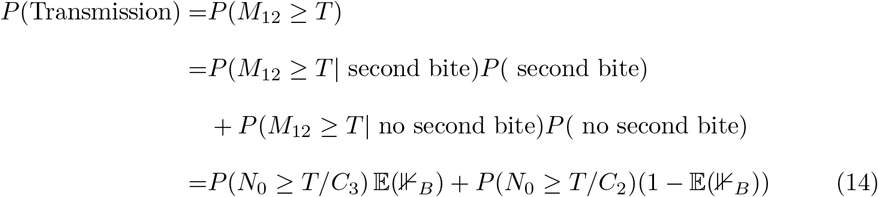

We will use Eq (14) to express how the variability in *N*_0_, which was the subject of interest in Doehl *et a*. [5], and the variability in the availability of a blood meal, which was the subject of interest in Serafim *et al*. [9], both contribute to the probability of disease transmission.

To help progress our arguments here we appeal to Chebyshev’s inequality, which tells us that a random variable takes values close to its expectation with high probability, more precisely it says that the probability of the random variable being further than *k >* 0 standard deviations from the expectation is smaller that *k*^−2^ i.e.

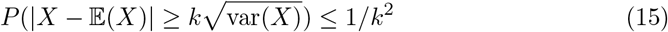

or equivalently

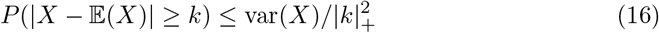

where we have introduced the rectifier function

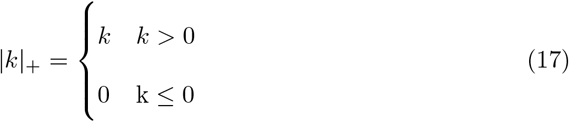

In order to accommodate negative *k*.

In the case when there is no bite at time *t* = 6 Chebyshev’s inequality allows us to put an upper bound on the transmission probability

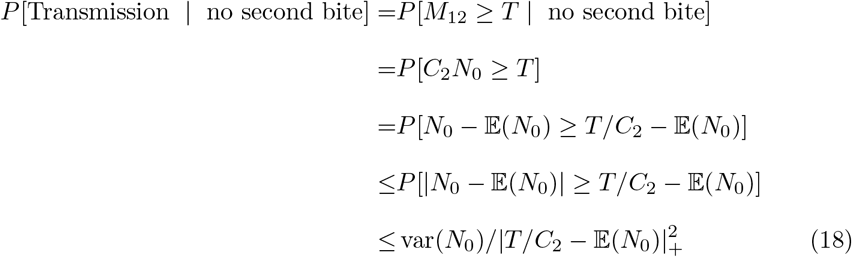

Such an upper bound is useful because it suggests ways the transmission probability can, in principle at least, be forced down. We could, for example, force down the variance of the number of parasites ingested at time *t* = 0. Alternatively, by decreasing the conversion rate from nectomonads at time *t* = 0 to metacyclics at time *t* = 12 we would decrease *C*_2_ which also serves to bring down the upper bound.

Considering the average over cases in which the blood meal bite does and does not occur at time *t* = 6, Chebyshev’s inequality leads us to an expression of the form

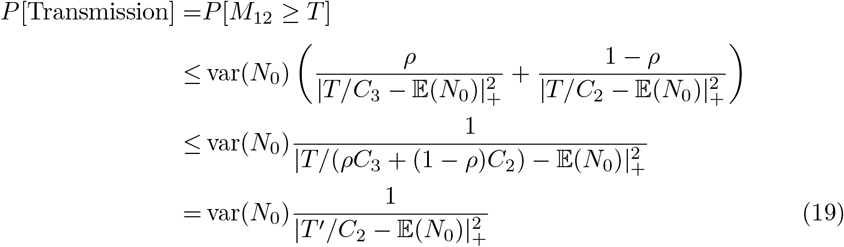

where the second line follows from Jensen’s inequality. Since *C*_3_ *> C*_2_, the second bite/retroleptomonad phenomenon effectively leads to a version of Eq (18) in which the transmission threshold has been lowered from *T* to

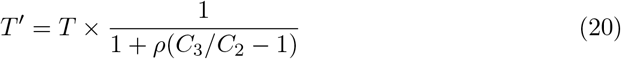

As well as providing quantitative predictions, this ‘equivalent threshold’ result is intended to provide another angle from which to interpret the significance of the retroleptomonad reproduction mechanism. Specifically, the retroleptomonads do not negate the capacity for skin heterogeneity to increase metacyclic numbers to transmission-sufficient levels for a subset of flies. Rather, they make these levels easier to attain. We see the effects of skin heterogeneity and the retroleptomonads act together to contribute to disease transmission.

An alternative expression linking the retroleptomonads to the transmission probability follows from assuming that the number of metacyclics derived from retroleptomonads is very large relative to the transmission threshold (i.e. *C*_3_*N*_0_ ≫ *T*). In this case we can consider the transmission probability given the blood-meal bite at *t* = 6 is close to one

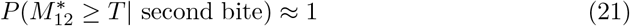

Then, using Chebyshev’s Inequality we see that

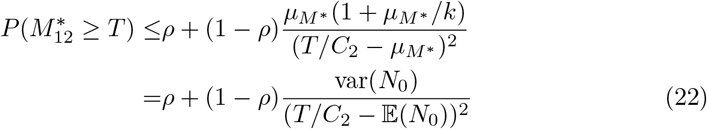

This bound provides another way to assess the relative influences of key parameters on the probability of transmission. For cases in which the transmission threshold is high relative to the number of metacyclics produced without the retroleptomonads (i.e. *C*_2_*N*_0_ ≪ *T*) and the blood-meal bite probability *ρ* is reasonable large, the rightmost summand in Eq (22) dominates. We then see the transmission probability reduced to the the blood-meal bite probability. When *ρ* is very small, however, the variance of *N*_0_, and the skin heterogeneity that drives it, becomes important again. It is this heterogeniety that provides each fly with its greatest chance of being able to deposit a sufficient number of parasites at time *t* = 12 to cause transmission. So in this case, where the blood-meal bite practically ensures transmission, we see that it is either the blood-meal probability or the skin heterogeneity that is most important for disease transmission.

### 1.4 Simulation study

This simple scenario is useful because it allows us to make analytical predictions about the behaviour of our system, however these predictions are useful only if we can verify that they apply to more sophisticated systems. Let us once more consider the full system for both models as originally defined (Model A: Eqs 1-3; Model B: Eqs 4-9). Each sexually mature female fly has a predetermined lifespan drawn from an exponential random variable of mean 13 days and bite throughout their lives, with inter-bite times drawn from a gamma distribution of mean 6 days and with bite loads as previously defined (Supplementary Method 2). We also reinstate the 3-day delay before the emergence of nectomonads.

We require a suitable metric to assess the infectiousness of *Leishmania* under a variety of P_B_ and k values. One such metric commonly used in epidemiology is the R_0_ [10] defined as “the number of secondary infections generated from a single infected individual introduced into a susceptible population” [11]. As we do not explicitly model individual hosts, this measure is unsuitable. Let us instead consider a proxy value: “Mean R_0_”, defined to be the average number of infections caused by a single sandfly. Though this is not strictly an R_0_ value, higher Mean R_0_ values imply a higher R_0_ value for the disease assuming that the number of sandflies biting a given infected host remains unchanged.

We determine that an transmission has occurred at a given bite using either a binary threshold or a smooth ‘threshold function’. In the case of the binary threshold, we assume that if the number of metacyclics transferred (M_T_) exceeds some fixed threshold T, we guarantee an infection (and if not an infection never occurs). For the smooth ‘threshold function’, we assume the chance of infection P_T_ at a given bite depends on M_T_ such that:

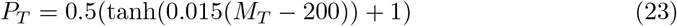

Whilst the binary threshold is easier to relate to our analytical work it is very unlikely to be applicable to a real situation, especially as it disregards any nutritional or genetic variation between potential hosts. Thus, let us consider the smooth threshold function. Corresponding figures for the binary threshold function can be found in the supplementary information, and we observe qualitatively similar behaviour with both the binary and smooth thresholds.

Let us first compare our two models for a range of different scenarios. Assume that some proportion of hosts is initially infected and that this proportion is fixed with no dependence on time or transmissions. Initially, we will consider two scenarios where either 100% or 25% of hosts are initially infected (for further scenarios see supplementary Figure 3):

**Fig 3.**
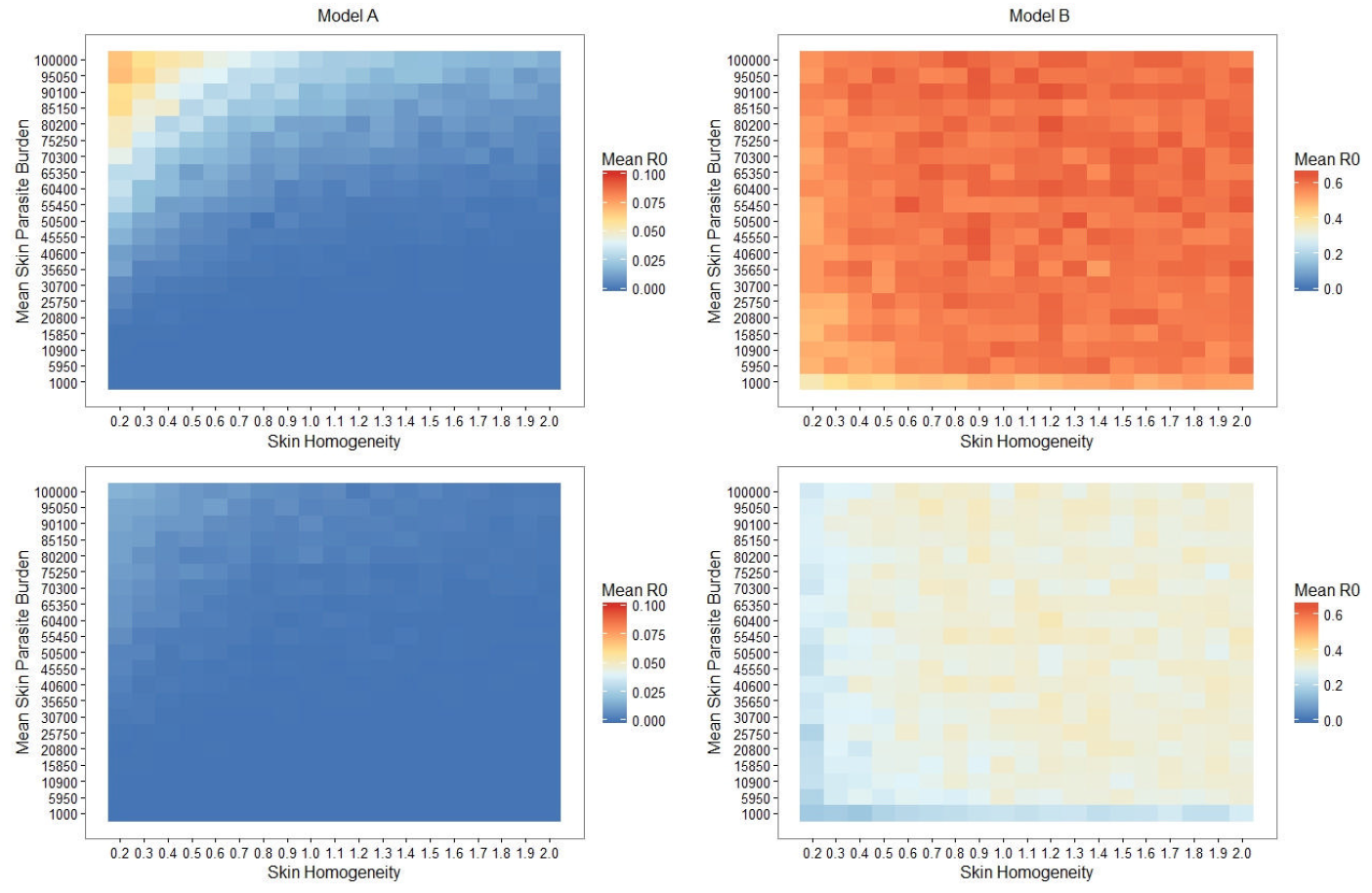
Retroleptomonad dynamics dominate over skin heterogeneity and result in elevated mean R_0_ values. Heatmaps of the Mean *R*_0_ for simulated sandflies for both Model A (left half) and B (right half) with 100% (top half) or 25% (bottom half) chance of biting an infected host. Note that each model utilises a different scale for clarity.

Although the simplest conclusion we can draw from these heatmaps is that introducing retroleptomonads increases our mean R_0_ value, there are a number of more notable results. We observe that for Model A there is a peak in the mean R_0_ value for low skin homogeneity and high mean skin parasite burden for both scenarios. Though our analytic approach does not deal directly with Model A, we could consider Model A to simply be the scenario where flies never take 3 bites (and thus where retroleptomonad lifecycle stage has no significant role). In this context we note that a low skin homogeneity increases the probability of transmission as some flies are now able to take up many parasites and remain infectious at the next bite, whereas for homogenous environments it is less likely that any fly would have this capability. This would match the prediction of Doehl *et al*. [5].

The peak is entirely absent from the corresponding heatmaps for Model B; instead we have a plateau spanning most of the parameter space with a slight decrease in mean R_0_ for very low k values (IE very patchy environments). We note from our analytical section that as *ρ* (the chance of taking 3 bites) increases, k (skin homogeneity) has a progressively reduced impact. Thus, given that *ρ* effectively remains constant (and non-zero) regardless of k one might anticipate that the mean R_0_ would be independent of k. Similarly, considering the magnitude of the amplification of the metacyclics (Fig 2A) it is reasonable to expect that the mean skin parasite burden would be relatively unimportant. This does not hold for very low skin homogeneity and/or parasite burdens, for under these conditions it is possible that the fly may fail to be initially infected or may not remain infected by the time of their second bite and thus be rendered unable to benefit from the retroleptomonad-dependent population boost.

Accordingly, skin homogeneity has a particularly reduced role in very long lived sandflies that bite many times. In such flies, the number of metacyclics are repeatedly amplified and this results in almost guaranteed transmission at the third and subsequent bites for the majority of these flies. To assess the impact of such flies, let us restrict the lifespans of the simulated flies to 20 days (see Fig 4A). Restricting the lifespan of the flies to 20 days appears to have minimal effect on the influence of skin homogeneity, though the plateau is now at a reduced mean R_0_ value (predominantly because flies cannot take as many bites). It should be noted that with a mean inter-bite time of 6 days, it is not unlikely that a given individual could take 3 bites in 20 days. Let us now consider a further restriction of the lifespan to 15 days (see Fig 4B).

**Fig 4.**
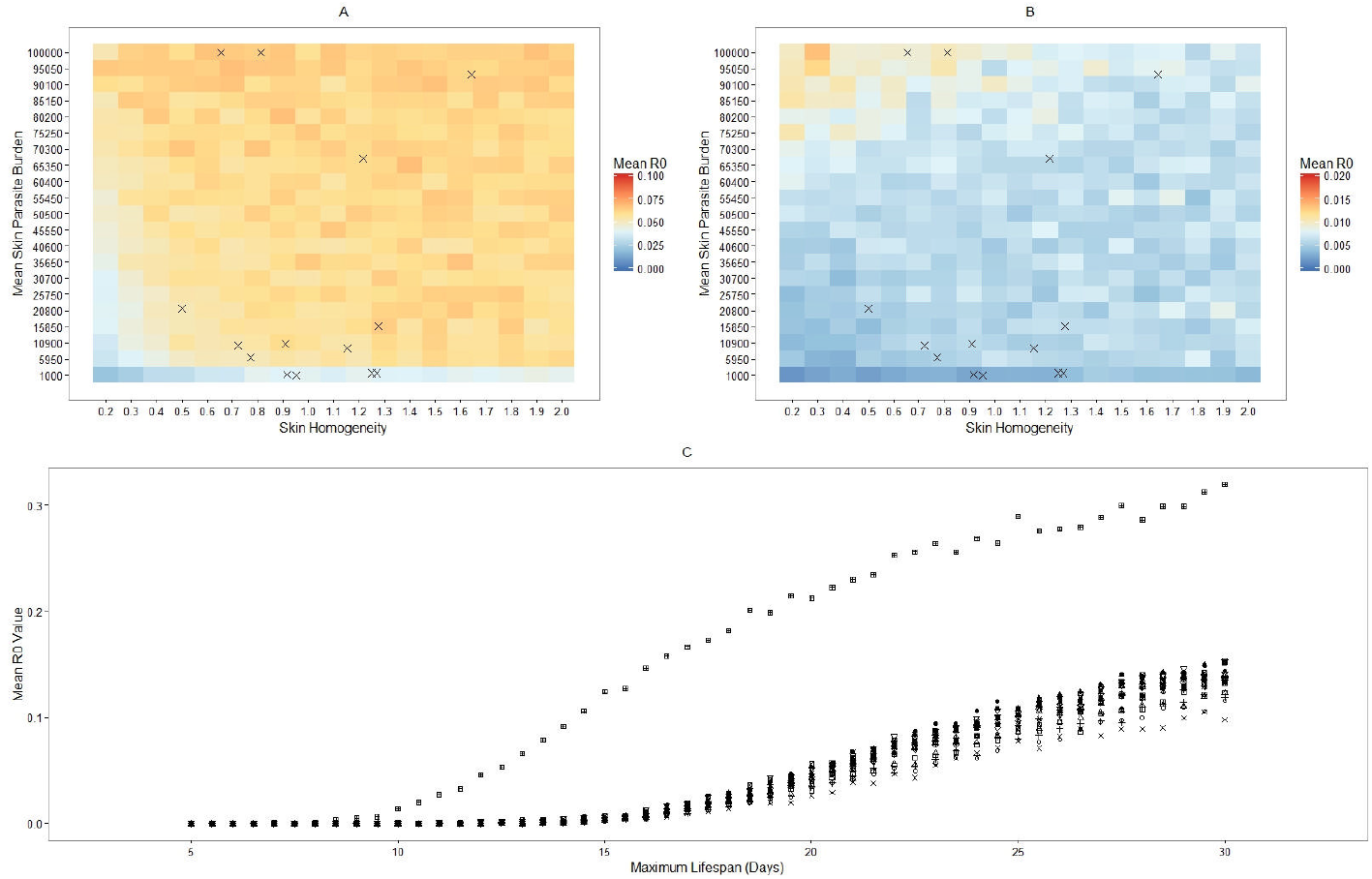
Retroleptomonad dominance is dependent on having a sufficiently large maximum lifespan. A, B) Heatmaps of the Mean *R*_0_ for simulated sandflies in Model B with 100% chance of biting an infected host and with lifespans restricted to 20 days (A) or 15 days (B). Crosses indicate the mean skin parasite burden and skin homogeneity of various mice from [5]. C) Mean R_0_ value against maximum lifespan for RAG mice 1-18 from Doehl *et al*.

Under this new, harsher restriction we see that skin homogeneity has much stronger influence on the mean R_0_ value. The peak observed in Model A is present again. The mean R_0_ value does not drop to zero away from that peak, however. This is likely because some flies will still manage to bite three times and thus benefit from the retroleptomonad replicative cycle (this could also be interpreted as having a low, but non-zero, *ρ* and thus we would expect a similarly low but non-zero mean R_0_).

Let us once more turn to the Doehl *et al*. mice to uncover the transition between these two states. Using the parameterisation for mice 1-18 from Doehl *et al* [5], we ran sets of 5000 sandflies for each mouse for a range of different maximum lifespans and calculated the mean R_0_ value for each set. We can then compare the trajectory taken by the mean R_0_ value for each population of simulated sandflies as we increase the maximum lifespan (Fig 4C).

We note that the mean R_0_ value increases with the maximum sandfly lifespan for all mice, especially once the maximum lifespan exceeds approximately 15 days, as anticipated from Figs 4A and 4B. This demonstrates the smooth transition away from a patchiness-dominated scenario and towards a retroleptomonad-dominated scenario as flies live for longer. Thus we see that although the conclusions of Doehl *et al* [5] do not hold for flies with unrestricted lifespans they may still be very much applicable to the shorter-lived portion of the population, and that reducing the maximum lifespan of the sandflies (and thus enlarging the shorter-lived portion) can have a tangible impact on the mean *R*_0_ value.

## Discussion

We observe both numerically and analytically that the inclusion of retroleptomonads allows sandflies which take multiple bites to transfer more parasites on subsequent bites and thus be more effective at transmitting leishmaniasis, as anticipated by Serafim *et al* [9]. Less trivially, we also observe that the inclusion of retroleptomonad-dependent amplification in the model alters the relationship between the mean R_0_ and skin homogeneity. In scenarios where the retroleptomonad life stage is absent (Model A) or play a substantially reduced role (Fig 4B) we see a strong dependence on skin homogeneity, with patchy environments leading to more transmissions as some flies take up many parasites and can then cause infections, as predicted by Doehl *et al* [5]. In scenarios where retroleptomonads are more important however, we see the opposite: skin homogeneity is unimportant to the transmission of the disease, as even small numbers of parasites initially present can be amplified greatly.

Although this result casts some doubt on the predictions made by Doehl *et al*. [5] there are other important considerations. Doehl *et al*. predicted that patchy skin distributions would enhance transmissions because flies could occasionally take up many parasites and then cause many infections, whereas in less patchy environments flies would be very unlikely to take up enough parasites to infect at the next bite. While we observe the loss of the relationship between skin homogeneity and mean R_0_ for the full system there are scenarios where it re-emerges. Flies with short lifespans (Fig 4B) cause more transmissions with patchy skin distributions than even ones. Such flies are unlikely to live long enough to bite three or more times and thus do not typically benefit from the inclusion of the retroleptomonad stage in the model. This is reflected in our analytical work. Consider the short-lifespan flies to have a low chance of taking three bites (IE a low *ρ*), then from Eq 22 we see that low k values increase the chance of transmission. Thus, there are conditions under which the scenario posed by Doehl *et al*. is relevant to the spread of the parasite.

The extent to which this applies in reality is uncertain. Although previous lab-based studies suggest that sandflies have fairly short adult lifespans (*<*20 days) [12] with further reductions when infected [13], release-recapture studies in natural settings suggest they may live much longer than in lab environments [14], especially given the additional mortality associated with oviposition in lab populations [15]. However, there is uncertainty as to the magnitude of the lifespan reduction in lab conditions, given that the lifespan of wild sandflies will vary between species and environmental conditions. Until this issue is clarified it will be difficult to properly quantitatively assess the importance of the retroleptomonad lifecycle stage in disease transmission.

This is further complicated by the feeding behaviour of the sandflies. We chose to model the time between subsequent bites (in days) using a gamma distribution of mean 6. Though this is a reasonable approximation for our model, in reality there is little information available about how often sandflies feed. It is likely that the feeding rate is linked to the oviposition cycle (given the dependence of oviposition on a blood meal) and the abundance of potential blood sources. Though the scenario of regular feeds posed by Serafim *et al* [9] is likely to be appropriate for a population with abundant sources of blood meals it is not necessarily true for all populations. Additionally, although we consider populations with different proportions of initially infected hosts (P_i_) including values such as 25% and 10% which are more applicable to current populations where it is endemic ([16, 17]) and observe that our results hold for such scenarios, we assume that hosts are evenly distributed throughout the populations and this is unlikely to be applicable in reality.

There is significant evidence that the behaviour of the sandflies is also altered once infected. A notable component of *Leishmania* infection known to alter sandfly behaviour is Promastigote Secretory Gel (PSG), a filamentous proteophosphoglycan-based gel secreted into the thoracic midgut and stomodeal valve [2, 3]. The occupation of the midgut by PSG causes the sandflies to feed ineffectively, taking smaller blood meals [3, 18] and demonstrating increased persistence when disturbed (with an increased likelihood of biting a second host after a disturbance) [13]. PSG also acts as a filter allowing only metacyclics to pass through [3], and impedes the unidirectional flow of blood through the stomodeal valve, causing the sandfly to regurgitate PSG and the parasites within it into the bite. This may amplify the number of infectious parasites transferred to a new host on a successful bite [3, 19]. These influences could have important implications. The frustrated feeding and increased persistence of infected sandflies suggests that PSG could increase the likelihood of transmission, both directly by transferring more infectious parasites and indirectly by increasing the rate at which retroleptomonads emerge. Although we model the regurgitation of parasites by increasing the number of transferred metacyclics for heavily infected flies [20] we do not directly model the PSG due to insufficient information regarding its production and how it interacts with the parasites in the midgut. Future studies may elucidate this matter further and allow a reasonable model to be produced.

Another avenue of future enquiry that holds potential value relates to improving the parameterisation of our model. As the discovery of the retroleptomonad lifecycle stage is very recent [9] we have insufficient data to properly parameterise Model B. Although we can demonstrate that our model produces similar behaviour to that of the experimental system, it would be preferable to have more data to base our parameter upon. Future studies may seek to improve the identification of retroleptomonads using transcriptomics tools as has been done for previous life cycle stages [21]. Alternatively, they may seek to provide more information about the two lifecycle stages we omit from our model, the amastigotes and procyclic promastigotes. Either of these options would greatly improve our model.

## Conclusion

This work has produced a basic population dynamic model for nectomonad, leptomonad and metacyclic promastigotes and integrated the recently discovered retroleptomonad promastigote. This model can be further enhanced via the addition of missing life cycle stages or additional parameter to improve the fit. This provides a basic tool that can be expanded upon depending on the aims of a study. For example, a similar model may prove useful if modelling the impact of interventions on promastigote dynamics. Through using Monte Carlo Simulations, we have demonstrated that the addition of retroleptomonads to the model greatly enhances transmission from the second bite onwards. This could suggest that retroleptomonads are a good stage to target in control efforts, potentially through interventions that reduce the number of bites a sand fly takes. We have also demonstrated that skin parasite heterogeneity does have an impact on *Leishmania* transmission, although a much smaller impact than retroleptomonads. A patchy distribution slightly enhances transmission when retroleptomonads aren’t present (such as the first bite), but a non-patchy distribution enhances transmission when retroleptomonads develop.

## Materials and methods

Model parameterisation was performed in RStudio v1.2.5019 (R version 3.6.1) with the digitize package [22] using data from [3] (see Supplementary Method S1 for full details). All Monte Carlo simulations were performed in MATLAB R2019b. Data analysis was performed in RStudio v1.2.5019 (R version 3.6.1).

## Supporting information

### S1 Method. Parameterisation of Model A

The basic model (Model A) produced in this study focuses on three promastigotes stages, nectomonads, leptomonads and metacyclics. Differential equation based models were produced based on the lifecycle described by Rogers *et al* [3]. This method assumes that the parameters used for rates are all constant. Parameterisation was achieved by fitting data from Rogers *et al* to the model. Data was collected from Figure 1 of this paper using the digitize function in R as this data wasn’t readily available [3, 22]. The digitize function is used to manually collected data from plots. The data collected were the total number of parasites in the sand fly and the percentage of each promastigote stage (nectomonad, leptomonad, metacyclic) present in the sand fly over a course of 10 days. This was then used to calculate the number of each promastigote stage. This data was then exported into MATLAB where the function “lsqcurvefit” was used to produce the best fitting parameter values. The quality of fit was assessed via an R^2^ value, which defines the proportion of variation that is explained by a model. A high R^2^ is indicative of a good fit where as a low R^2^ is indicative of a poor fit.

### S2 Method. Bite Mechanics

In order to represent a ‘patchy’ environment for sandflies to draw parasites from, bite loads were generated from a negative binomial distribution using the ‘nbinrnd’ function in MATLAB. This function outputs a random value from a negative binomial distribution. This takes the following inputs: P (Probability of a positive result, in this case probability that a sand fly will ingest parasites following a bite) and R (the number of successes required). R and P are defined as follows:

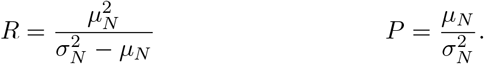

where:

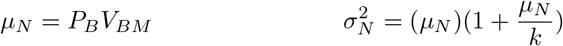

Since this model starts from nectomonads, the number of amastigotes had to be converted to nectomonads. The number of nectomonads is approximately three times greater than the number of amastigotes [3], hence the number of amastigotes was multiplied by three to estimate the number of nectomonads.

**Fig 5. Supplementary Figure 1:**
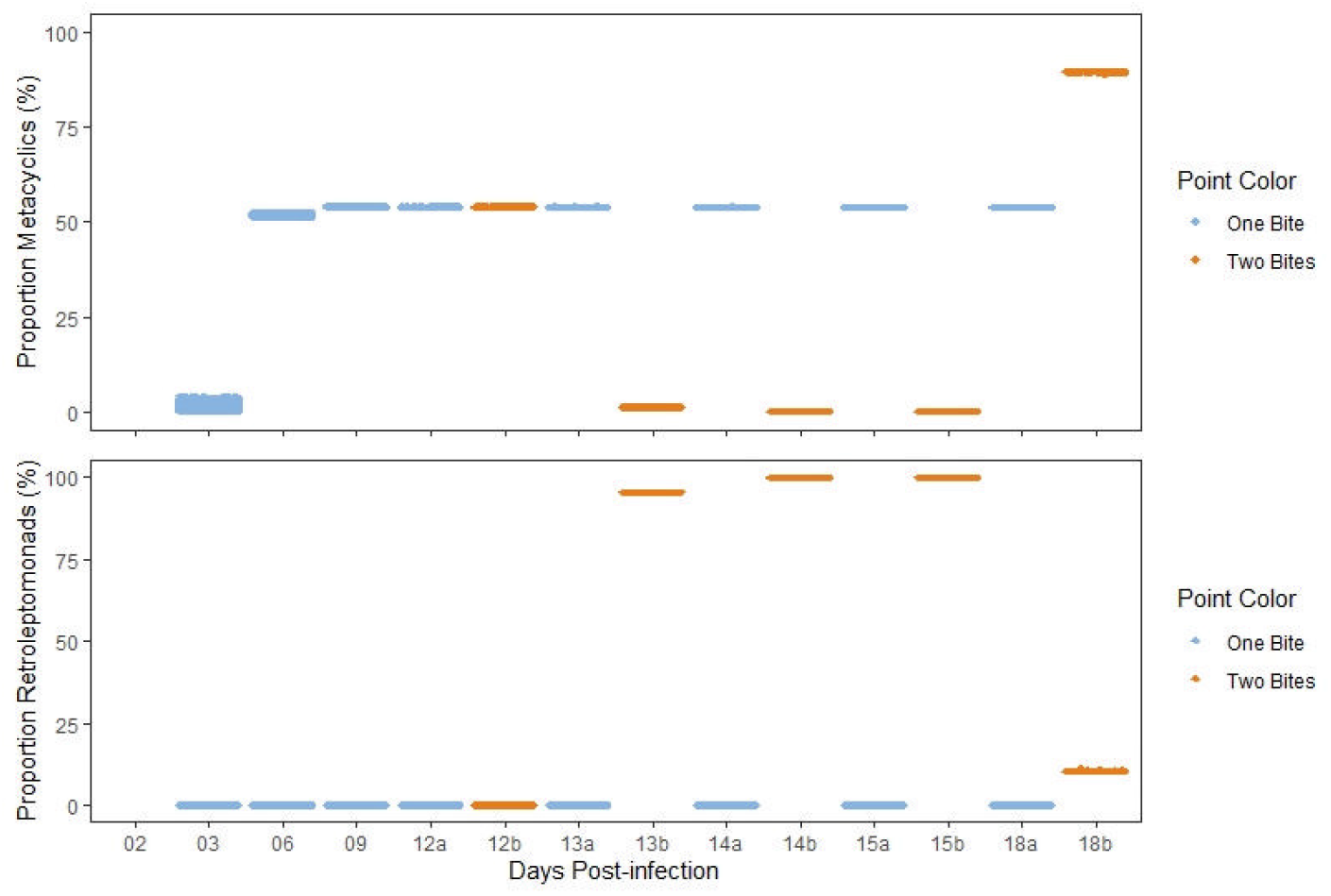
**Replicating the results of [9] (parasite proportions).** Comparison of the proportions of metacyclics (top) and retroleptomonads (bottom) at specific days throughout the lifespan of the simulated flies. Blue represents flies that bite only at day 0, orange represents flies that bite at day 12. The two categories are combined prior to day 12.

**Fig 6. Supplementary Figure 2:**
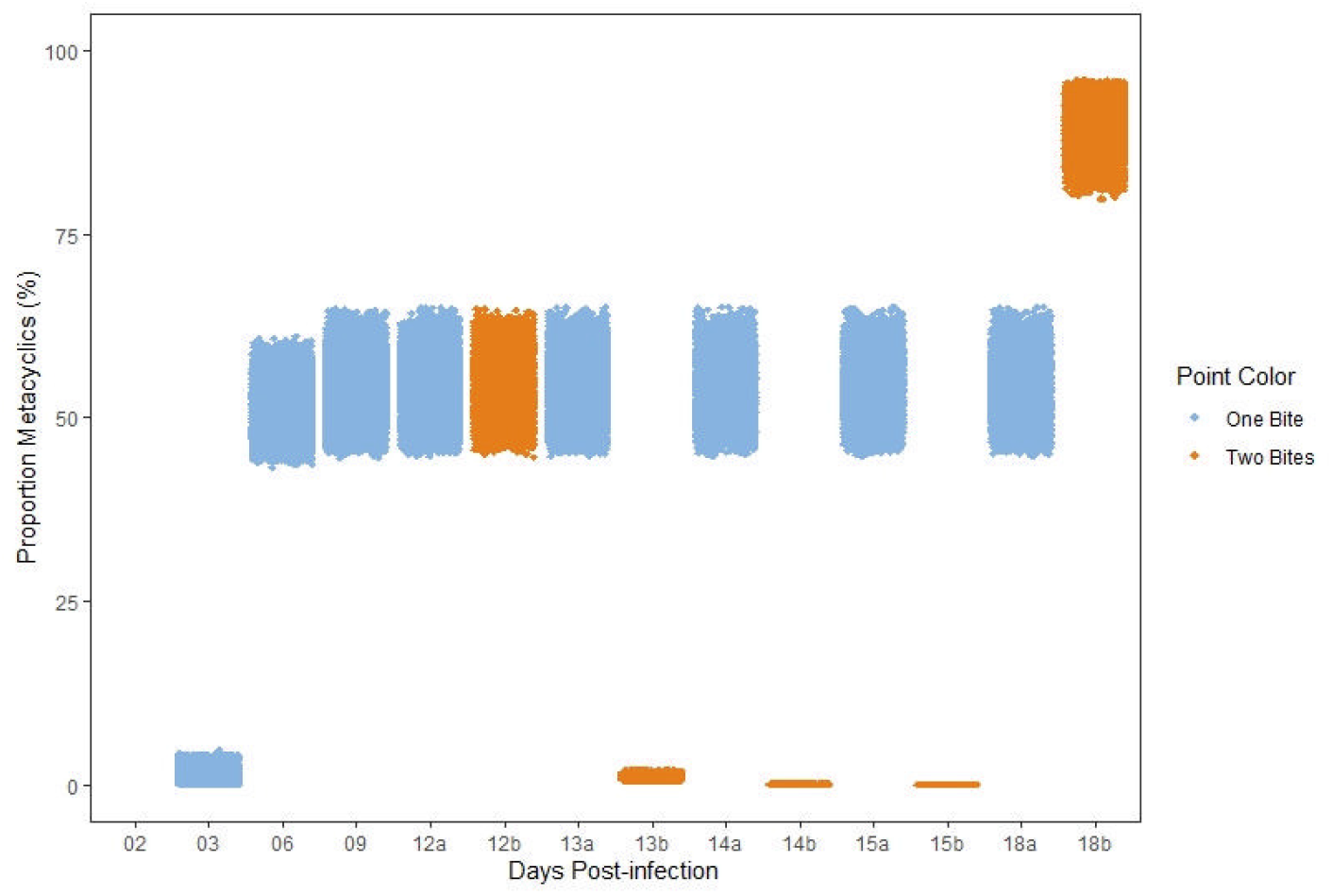
Evaluating model robustness by randomising parameters. Number of metacyclics within the sandflies at specific days, with all parameters randomised prior to the start of each simulation. Parameters lie within 10% of the default value (Table 1). Blue represents flies that bite only a day 0, orange represents flies that bite at day 12.

**Fig 7. Supplementary Figure 3:**
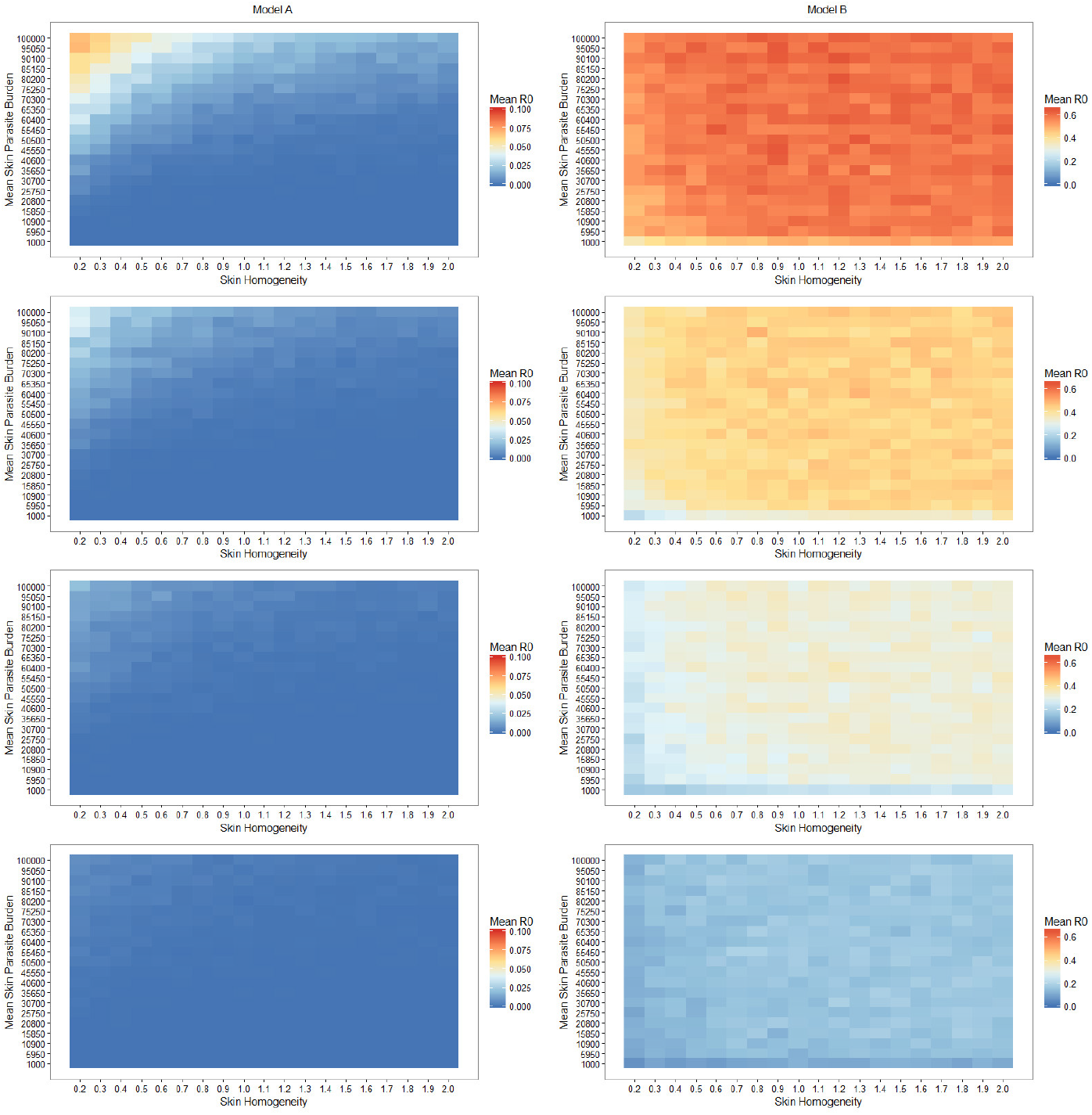
Additional infected host proportions reflect the retroleptomonad dominance. Heatmaps of the Mean R_0_ for simulated sandflies for both Model A (left half) and B (right half) with 100% (top row), 50% (second row), 25% (third row), and 10% (bottom row) chance of biting an infected host, with the smooth transmission threshold function.

**Fig 8. Supplementary Figure 4:**
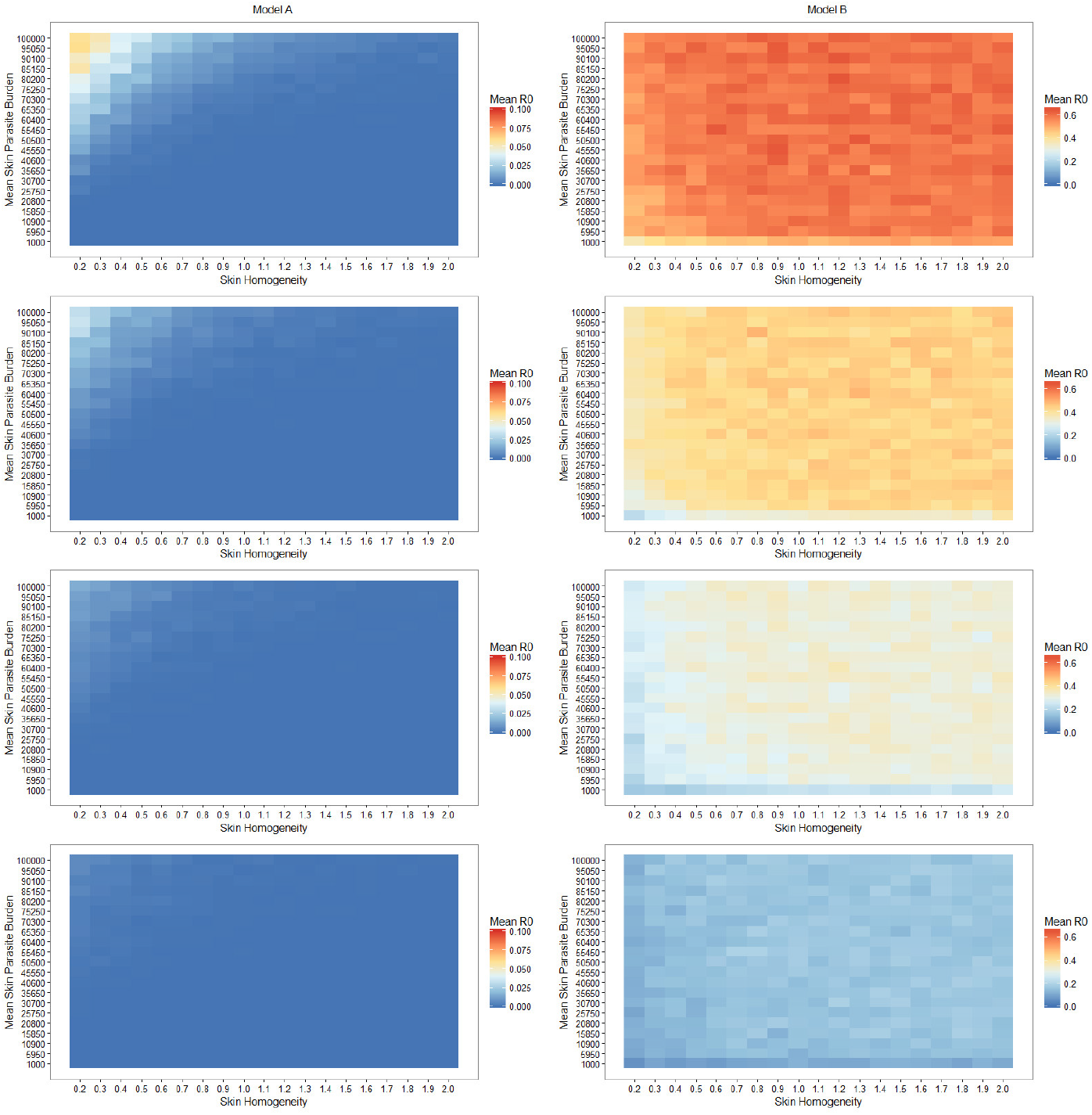
Heatmap dynamics remain qualitatively similar under a binary transmission threshold. Heatmaps of the Mean R_0_ for simulated sandflies for both Model A (left half) and B (right half) with 100% (top row), 50% (second row), 25% (third row), and 10% (bottom row) chance of biting an infected host, with the binary transmission threshold.

**Fig 9. Supplementary Figure 5:**
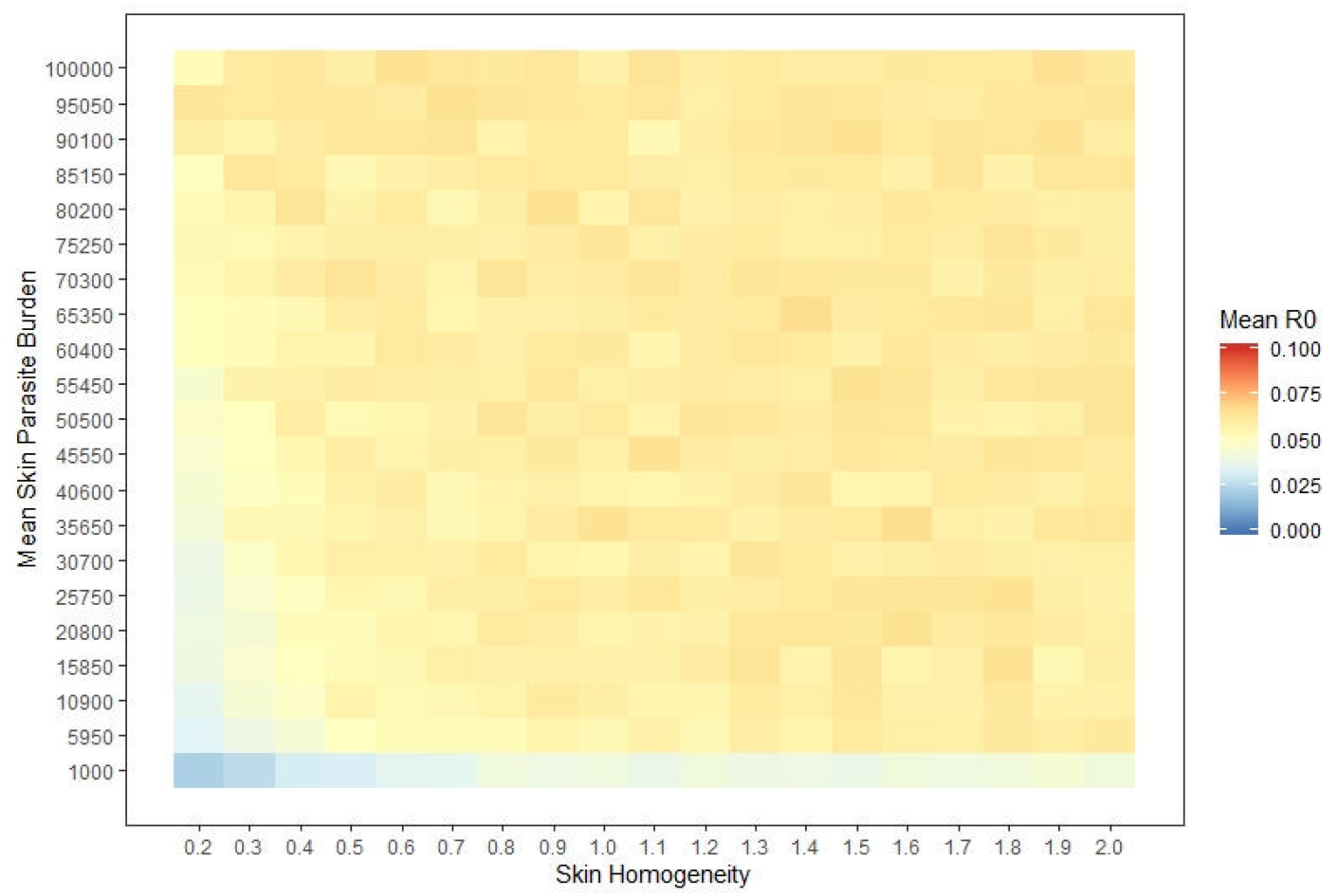
Reduced lifespan (20 days) dynamics remain qualitatively similar under a binary transmission threshold. Heatmap of the Mean R_0_ for simulated sandflies in the retroleptomonad model with 100% chance of biting an infected host and with lifespans restricted to 20 days, with the binary transmission threshold.

**Fig 10. Supplementary Figure 6:**
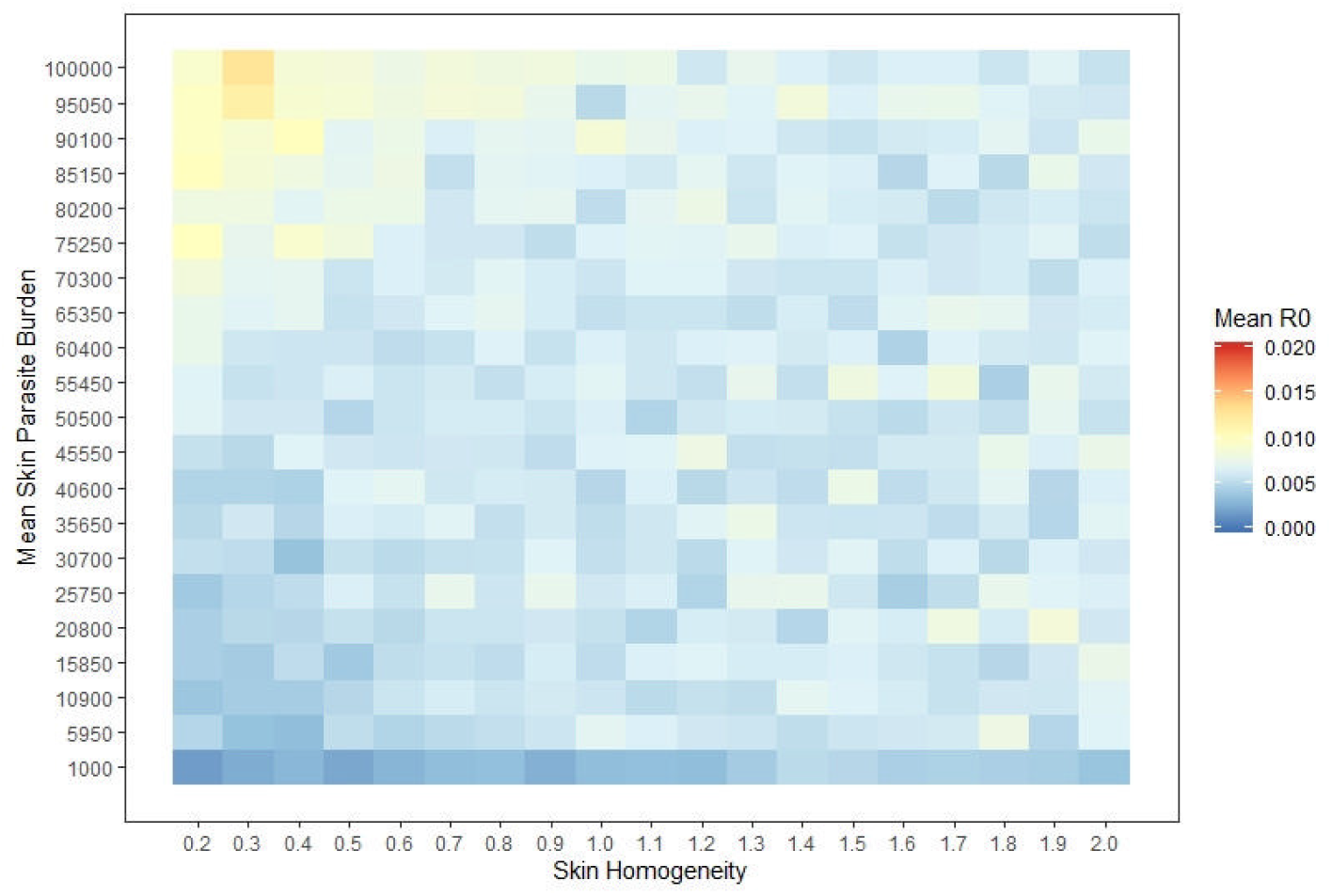
Reduced lifespan (15 days) dynamics remain qualitatively similar under a binary transmission threshold. Heatmap of the Mean R_0_ for simulated sandflies in the Retroleptomonad model with 100% chance of biting an infected host and with lifespans restricted to 15 days, with the binary transmission threshold.

## Notes

### Competing Interest Statement

The authors have declared no competing interest.

